# Understanding science communication in human genetics using text mining

**DOI:** 10.1101/2020.07.24.219683

**Authors:** José J. Morosoli, Lucía Colodro-Conde, Fiona K. Barlow, Sarah E. Medland

**Author notes:** Correspondence and requests for materials should be addressed to J.J.M.

## Abstract

We conducted the first systematic text mining review of online media coverage of genome-wide association studies (GWAS) and analyzed trends in media coverage, readability, themes, and mentions of ethical, legal, and social issues (ELSI). Over 5,000 online news articles published from 2005 to 2018 all over the world were included in analyses. Our results show that while some GWAS attract a great deal of online interest many are not reported on, and that those that *are* covered are described in language too complex to be understood by the general public. Ethical issues are largely unaddressed, while suggestions for translation are increasing over time. Our review identifies areas that need to improve to increase the effectiveness and accuracy of the communication of genetic research findings in online media. We have also developed a website where all results described below can be explored interactively: https://jjmorosoli.shinyapps.io/newas/.

---

Over the last few decades, online news and media have become the main source of scientific information for many individuals and decision-makers^1^. Given the potential for media to set the public agenda, there is a need for science to understand and track how science is covered in the media^2^. Moreover, understanding how science is communicated is especially relevant in areas that can have marked social implications and divide public opinion, such as human genetics^3^. However, human genetics research has not engaged as strongly with an evidence-based, early approach to communicating communication, as other fields have (e.g. climate change, vaccines, stem cells, and genetically modified organism, etc.)^2,4^. Thus, the main goal of our review is to understand the portrayal of complex trait genetics in online media; additionally, we aim to demonstrate how big data analytics can inform science communication.

We retrieved PubMed identifiers (PMIDs) and citation metadata from all GWAS publications indexed by the NHGRI-EBI GWAS Catalog on 17 September 2018^5^. We classified the traits analyzed in these GWAS into non-disease and disease traits. Disease traits were further classified using the International Classification of Diseases 10^th^ Revision (ICD-10)^6^. We used Altmetric Explorer API^7^ to identify online mentions of GWAS publications (i.e., news sites and blogs) and retrieve metadata and URLs. Our analyses included 3,555 GWAS publications on 1,943 different traits, featured in 5,505 English language online news sites (see Fig. 1). Information about the GWAS publications and online references reviewed can be found in Supplementary Tables 1-5.

**Table 1.** Top 10 topics in the GWAS news corpus, labels assigned based on top terms within topic.

| **<u>Topic</u>** | **<u>Top 7 terms that contribute to each topic</u>** |
| --- | --- |
| Major Depression | Depression, disorder, scientists, university, condition, psychiatric, major |
| Cancer in women | Cancer, breast, lung, women, ovarian, cancers, common |
| Methods | SNP, association, loci, wide, analysis, studies |
| Asthma and empathy | Women, empathy, twins, birth, children, asthma, age |
| Educational attainment | Education, intelligence, differences, social, attainment, twins, environment |
| Facial genetics | Hair, skin, nose, facial, color, shape, pigmentation |
| Ancestry | African, American, ancestry, European, populations, variant, children |
| Alzheimer's disease | Brain, Alzheimer's, cognitive, memory, dementia, found, scientists |
| Diabetes | Diabetes, obesity, type, life, lifespan, health, diseases |
| Cardiovascular disease | Heart, blood, stroke, pressure, cholesterol, cardiovascular, coronary |

**Fig. 1.**
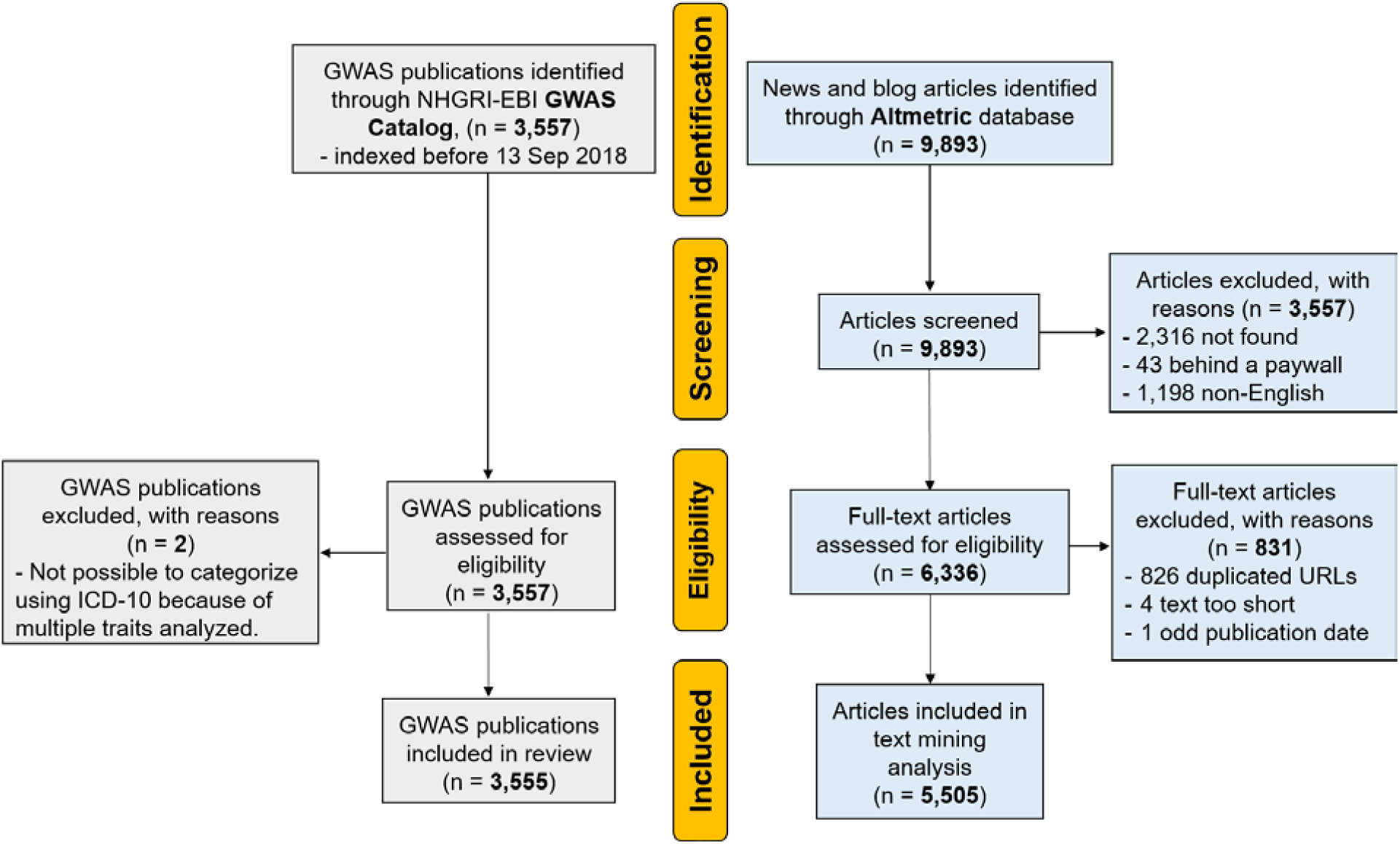
Systematic review flow diagram. Duplicated URLs refer to news sites and blogs that linked to more than one GWAS publication. In those cases, we only analyzed the website once. In the case of identical or almost identical websites (identical, aggregated, or rephrased), we analyzed all entries.

First, only 22.9% of published GWAS were covered online. Almost 40% of retrieved news and blogs articles contained identical, aggregated, or rephrased content from another publication. Both GWAS publications and their online coverage increased each year (see Fig. 2). The most frequently studied traits have been non-disease traits (33.7%), neoplasms (13.0%), and mental and behavioral disorders (10.4%). These were also the traits that most likely appeared on the news, receiving 43.4%, 11.6%, and 14.1% of all news reports, respectively.

**Fig. 2.**
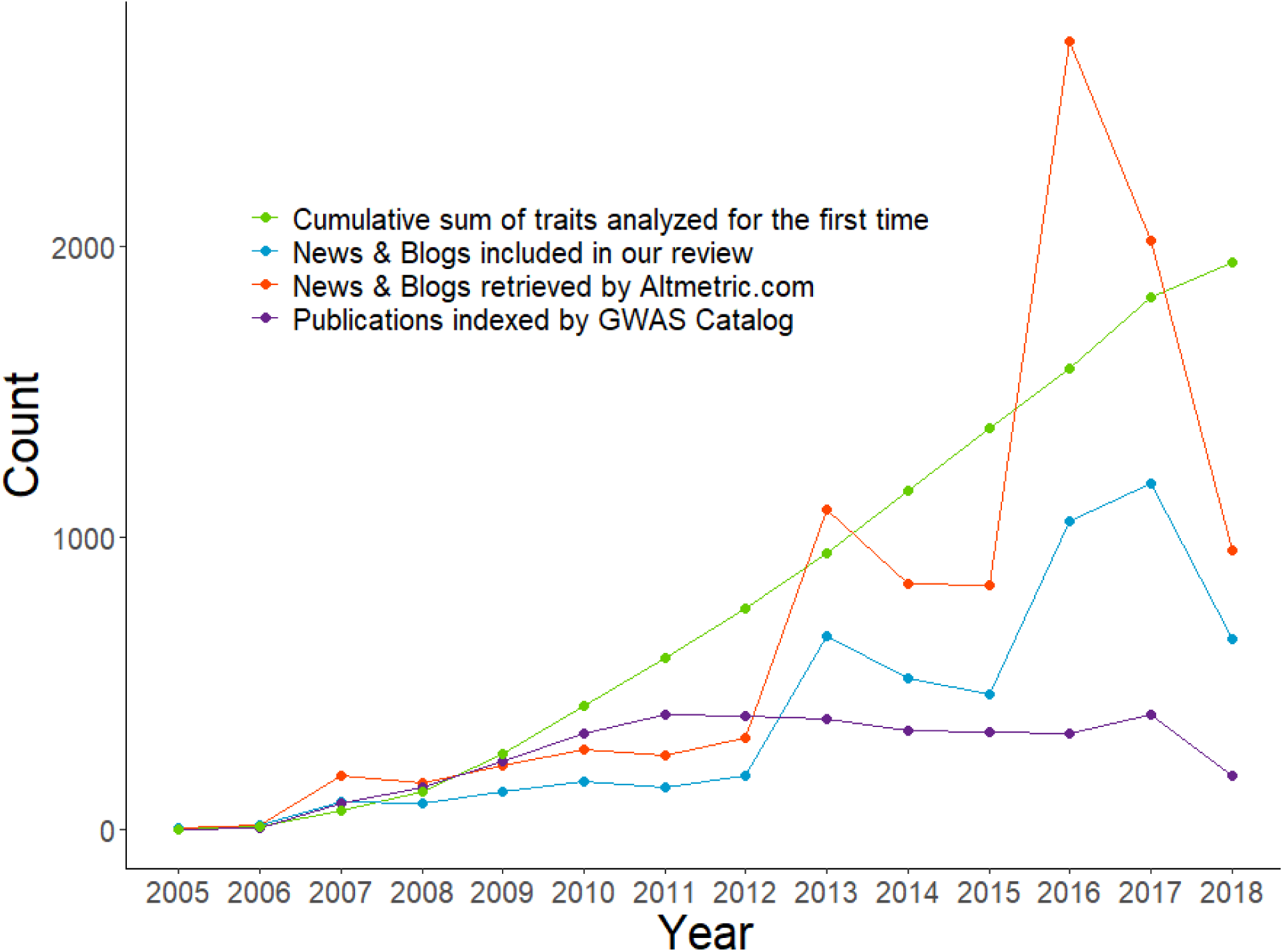
Number of GWAS publications and mentions in news and blogs by year. Online attention to GWAS has increased over time independently from the increase of GWAS publications per year (partial r = 0.69 [0.23,0.90]).

Having established rates of media coverage, we addressed the following research questions: 1) Is news coverage about GWAS understandable to readers?; 2) How technical is the news coverage about GWAS?; 3) How is genetic research being framed in the news?; 4) To what extent do news coverage address applications of genetic research as well as ethical, legal, and social issues (ELSI); 5) Why do some scientific publications receive more media attention than others?; and 6) What are the most common themes in the media?

A prerequisite for effective science communication in mass media is that the communication is understandable or readable by the intended audience. *Readability* refers to how easy to understand a text is. It depends on the content, style, design, and organization with prior knowledge, reading skill, interest, and motivation of the intended audience^8^, and can be measured using *readability formulas*. These formulas use vocabulary range and sentence length to predict text difficulty and estimate the level of reading skill required to understand it^8^. In order to provide a baseline of language complexity of online media, we calculated four readability indexes for each news site and blog: the Flesch Reading Ease score (FRES), Flesch-Kincaid Grade Level (F-K), Gunning Fog Index (GFO), and Simple Measure of Gobbledygook (SMOG). Across readability indexes, analyses showed that 95% of the news sites and blogs would require a college graduate reading ability. Placing this in a US context, about 46.3% of US adults hold an associate’s degree or higher, and general online media such as that produced by The Huffington Post or CNN is approximately five times less complex^9^. Moreover, guidelines for effective communication suggest aiming for two to five grades lower than the highest average grade level of your intended audience^10^, meaning that online coverage of GWAS is effectively inaccessible for approximately 64% of US adults^2^.

## Technical vocabulary

Another potential barrier to effective communication is *technical vocabulary*. In 2018, only 6.3% of those with a bachelor’s degree in the US majored in biological, agricultural, or environmental sciences – degrees that might feasibly introduce people to genetic terminology^11^. In order to estimate prevalence of genetic jargon in media coverage of GWAS, we calculated how many of the terms from the NHGRI Talking Genetics Glossary^12^ were present in each article. The 5 most common genetic terms across all websites were RNA (which was present in 65.3% of websites), risk (63.7%), gene (62.0%), genome (61.1%), and DNA (45.0%). On average, each website used 9 out of 231 terms present in the NHGRI Talking Genetics Glossary (M = 8.8, SD = 4.5). Note that ‘risk’ was used in 63.7% of news articles versus ‘susceptibility’ (12.2%) or ‘protect’ (11.3%); and ‘gene’ was used in 62.0% while ‘marker, ‘polymorphism’, or ‘allele’, where used in 15.9%, 11.6%, and 6.3% respectively. Core terms in complex trait genetics, such as ‘polygenic’ and ‘interaction’, only appeared in 2.9% and 6.7% of all news articles, respectively. The low use of genetic jargon may reflect an effort to make genetic research more accessible; however, the challenge lies on finding the right balance between using a more readable language while introducing contemporary genetic terminology. That is, eliminating jargon may not lead to more accessible science communication accessible and can remove the opportunity to explain more complex genetic concepts to readers.

## Framing

The interpretation of information can be influenced by the presence or absence of certain words, phrases, or images in an article through a mechanism known as *framing*^2^. Effective science communication involves the use of framing in a way that overcomes audience heuristics and personal motives that interfere with an accurate understanding of scientific knowledge^13,14^. Previous studies have argued that portrayal of genetics in the news has been simplistic, disregarded failures to replicate findings, promoted the nature *vs* nurture dichotomy, and ignored ethical callenges^15-18^, while finding heterogeneous levels of genetic determinism^18-20^. To identify the frames used in the news articles, we analyzed the use of key terms using an adaption of the framework developed by Carver and colleagues^21^ which classifies representations of genetics across traits and time as: materialistic, deterministic, relativistic, evolutionary, or symbolic (see Box 1).

### Box 1

**Gene Framing Scheme, modified from Carver**, et al. ^22^. Asterisks indicate that all derived words from that root word were included.

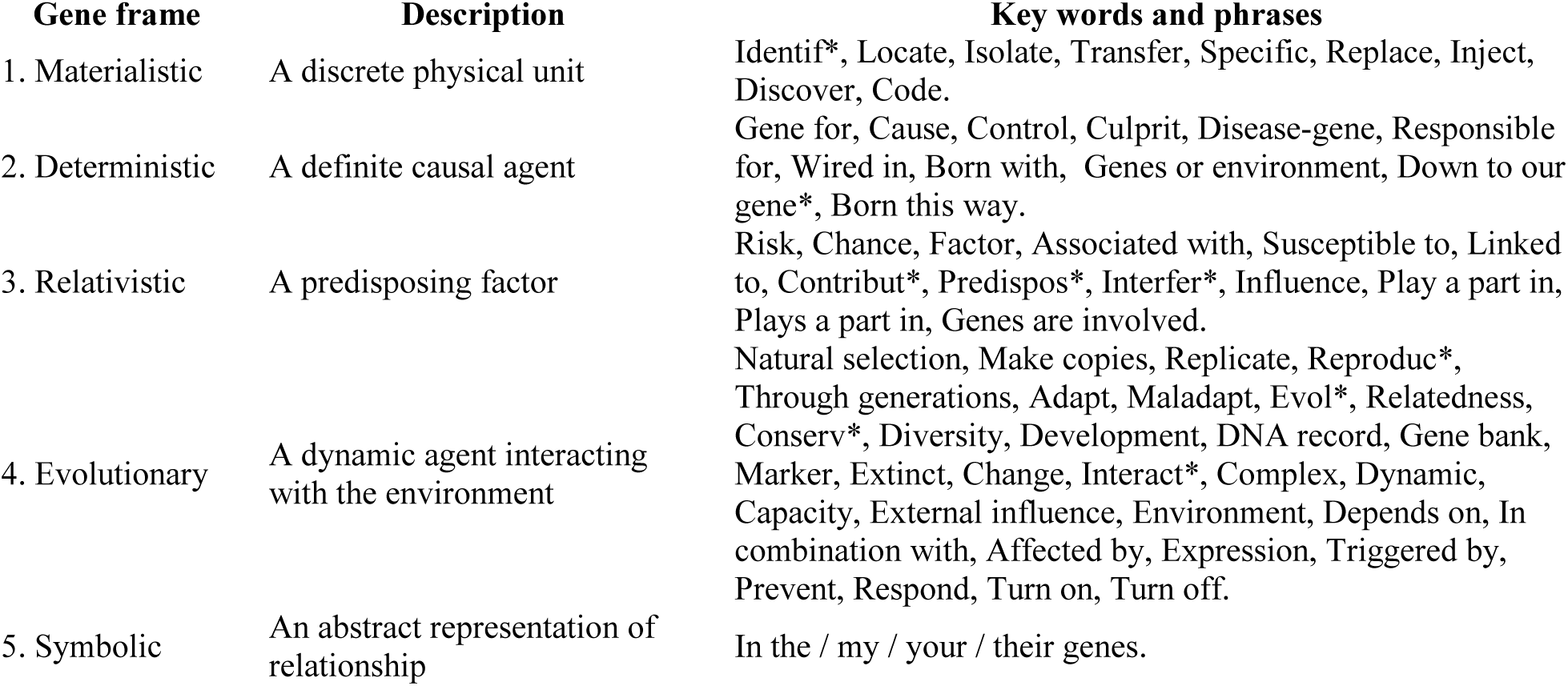

Relativistic and materialistic frames were the most frequent frames used in news sites and blogs. Use of deterministic key words and phrases was comparatively lower. This pattern is largely stable across time, although in mental and behavioral disorders we observe a decrease in use of deterministic terms and an increase of relativistic terms (see Fig. 3 and our website^23^ for more details). In comparison with a previous review analyzing news content about genetics published in tabloid and elite newspapers in 2005-2008^20^, we found the same average usage of deterministic (16.2%) and evolutionary frames (12% *vs* 12.9%), and higher average usage of relativistic (35.8% *vs* 13.5%) and materialistic frames (34.3% *vs* 25.6%). Notably, the phrase ‘*gene for*’ was rarely used between 2005 and 2012, but appeared more frequently from 2013^*2*3^. As previous work has found that genetic explanations of human behavior often activate deterministic asummptions^24,25^, despite the relatively low presence of deterministic words, a single deterministic catchphrase might override complex explanations of genetics, especially given the low readability of news coverage on GWAS.

**Fig. 3.**
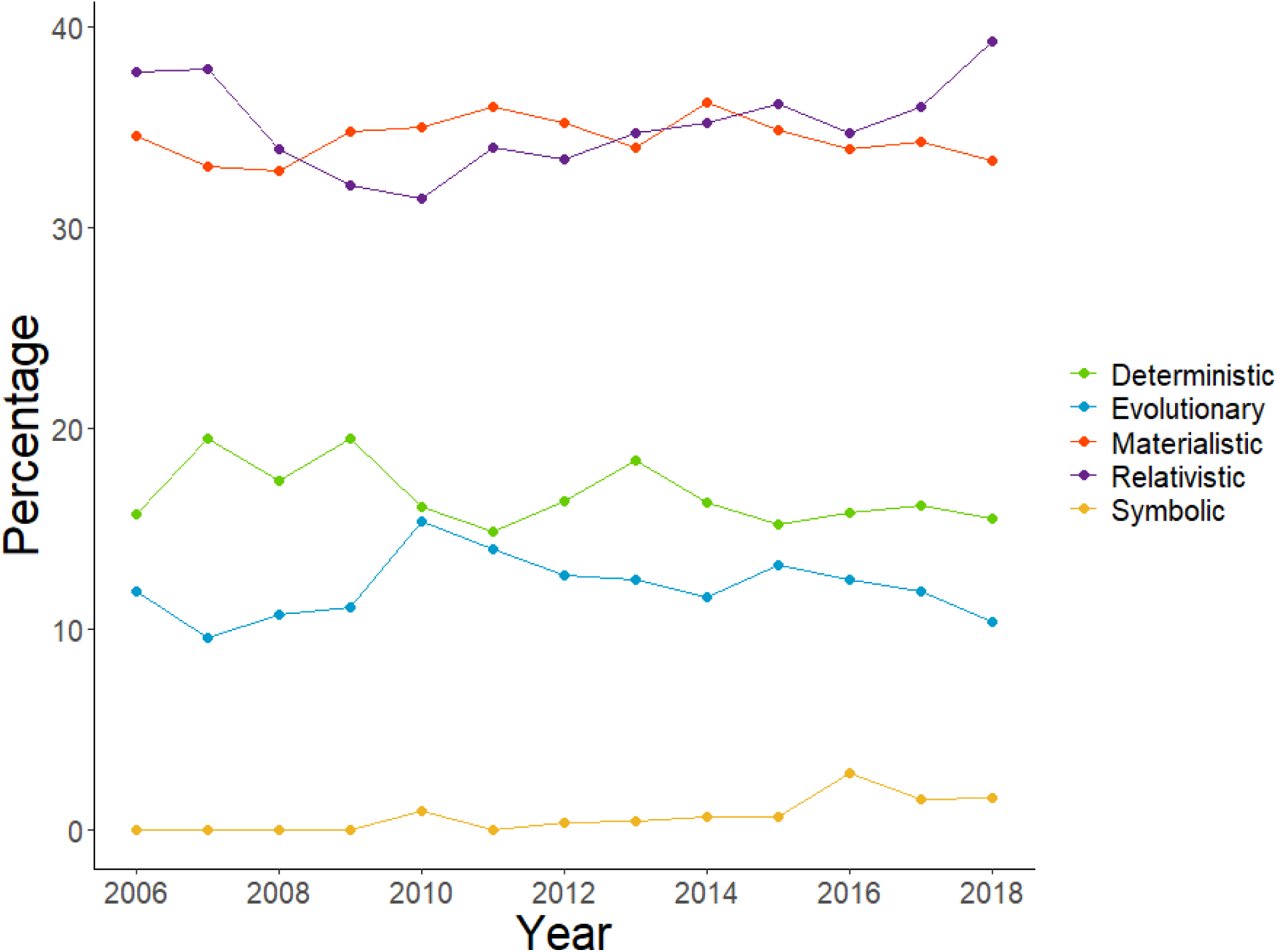
Relative use of frames over time across all traits, data points indicate average presence of frame within that year across all news sites and blogs.

## Implications of Genetic Findings

Concerns about the *implications* of genetic findings, regarding privacy, insurance coverage, and discrimination are widespread^26^. Therefore, we analyzed the frequency of use of terms associated with: a) translation of genetic research; b) ELSI; and c) positive and negative emotions; within news coverage, across time and traits (see Supplementary Table 5).

We found that mentions of clinical implications in GWAS coverage have increased over time. The terms ‘prevent’ (24.8% of 5,505 news articles), ‘therap*’ (23.7%), ‘screening’ (7.4%), ‘precision medicine’ (2.4%), ‘detection’ (2.3%), and ‘pharmacogenomics’ (0.9%), all started to appear more frequently from 2015 onwards^23^. However, there was relatively little change in the use of the terms related to ELSI, such as ‘policy’ (2.7% of news articles), ‘ethic*’ (2.1%), ‘minorit*’ (1.9%), ‘privacy’ (1.4%), ‘stigma’ (1.2%), ‘discrimination’ (0.9%), ‘insurance’ (0.7%), and ‘eugenic*’ (0.9%), which remained low over the years^23^. Given public concerns about privacy and potential discrimination^26^, researchers are encouraged to consider mentioning ELSI processes when reporting genetic findings, and considering early public engagement when working on traits that have the potential to become contentious^2^. Finally, positive words were used more frequently than negative words, although the prevalence of both positive and negative terms was low, suggesting that news coverage was not typically emotionally valenced. The most common positive adjectives were ‘novel’ (used in 14.7% of news articles), ‘unique’ (6.8%), and ‘robust’ (4.4%), while most common negative adjectives were ‘weak’ (7.6%), ‘ineffective’ (0.8%), and ‘inadequate’ (0.7%).

## Predicting news coverage

Next, we assessed the claim that science journalism largely relies on measures of relevance provided by science itself to choose *which publications to cover*^1^. Using the metadata from the GWAS Catalog and the Journal Citation Reports^27^ we used negative binomial regression analyses to establish if the number of online mentions of GWAS publications were predicted by year of publication, number of significant loci, discovery sample size, or journal impact factor. Publications that were more recent (Incidence rate ratio (IRR) = 1.88 [1.71, 2.06], P < 0.001), with bigger sample sizes (IRR = 2.34 [1.83, 3.05], P < 0.001), and published in journals with higher impact factors (IRR = 2.67 [2.39, 3.01], P<.001), received more media attention^23^. There were some significant interactions with year of publication: in later years sample size was less predictive of coverage (IRR=0.69 [0.57, 0.83], P < 0.001) while impact factor was more predictive (IRR = 1.12 [1.02, 1.23], P = 0.015). There was also a significant interaction between impact factor and sample size (IRR = 0.87 [0.77, 0.97], P = 0.003). This regression model explains 38.7% of the variance and supports the notion that a measure of relevance, such as impact factor, influences which stories get covered, and that this has become more salient in recent years. Examination of residuals showed that while the model was accurate for most traits, there were substantially more online mentions than predicted by our model to studies on neoplasms, behavioral disorders, chronotypes, intelligence and educational attainment, and alcohol and coffee consumption, suggesting differential trends in media interest. Comprehensive results from the regression analyses are available in our website.

## Themes

Finally, we conducted a *topic model analysis* in order to identify overarching themes in news coverage. Topic modeling classifies words into natural categories based on their co-occurrence within a document. In the online news analyzed, a model with 30 topics showed the best fit. The top five topics in the news coverage were major depression, cancer in women, asthma and empathy^1^ and educational attainment (see Table 1). We classified each article based on the topic it most likely belonged to, which allowed us to explore the context in which the key words and frames described above were used. The topic ‘sleep disorders’ had the highest use of deterministic key words and ‘major depression’ had the lowest. The word ‘environment’ was most frequently used when talking about ‘immune system’ and ‘educational attainment’, while ‘eugenic’ was most frequently used with the topics ‘educational attainment’ and ‘cancer in women’.

In summary, in the first systematic text mining review of online media coverage of GWAS research we characterize the use of technical vocabulary, themes, emotional valence, and topics, among others. We also identify some areas that we as a scientific community need to improve upon: online media coverage of GWAS should be written so it is more accessible, include more modern genetics terms, and more consideration of ELSI issues. We argue that science communication research in our field can benefit from big data and text mining techniques which can identify, like the present review, areas in which we can improve our communication practices in order to adapt to the evolving online media landscape.

## Methods

Published GWAS and associated metadata were retrieved from the GWAS Catalog^5^. Online mentions of publications were obtained via a research agreement with Altmetric^7^. Altmetric tracks mentions of papers in the news by i) searching for a direct hyperlink to a scholarly paper in the content of a news report, and ii) searching the news report’s text for mentions of the scholarly paper, journal, and author(s). More details can be found in: https://www.altmetric.com/about-our-data/our-sources/news/. Each URL was accessed and coded as found or not found and reasons were reported (see Supplementary Table 4). Data analysis was conducted in R-3.6.2^28^. Text in the online mentions was retrieved by hand, stored as text file, and analyzed using tidytext^29^. The interactive website was developed using shiny^30^.

Structural Topic Modelling as implemented in the stm package^31^ was used to identify latent topics in our text corpus. The optimum number of latent topics was decided based on i) highest held-out likelihood and semantic coherence and lowest residuals, which lead us to choose a 30 topics solution^32^.

For the regression analysis, the dependent variable ‘online mentions’ was based on the number of online mentions originally identified by Altmetric, (including news articles in languages other than English for which we were not able to retrieve the text; see Fig.1). Journal impact factor for GWAS publications was defined as impact factor in the year before the paper was published. The distribution of the dependent variable ‘online mentions’ was highly skewed and a negative binomial regression model was preferred over a linear model. Regression analysis was conducted using glmmTMB^33^.

Relative usage of frames in an article was computed by dividing the proportion of key terms of a specific frame by the weighted sum of total key terms of all frames present in that article within article. First, we computed the presence or absence of each of the key terms associated with the five possible frames within each article. Then, we calculated the relative usage of each frame (see formula below).

|  |  |
| --- | --- |
| $\frac{\frac{n_{i,j}}{m_j}}{\sum_{j=1}^5 \frac{n_i}{m_j}} \cdot 100$ | $n$: number of frame $j$ keywords present in news article $i$<br>$m$: maximum possible number of keywords for frame $j$ |
| **Example**<br>News article #614<br>$n_{614,materialistic} = 1$ (out of 13); $n_{614,mat}/m_{mat} = 1/13 = .077$<br>$n_{614,deterministic} = 0$ (out of 11); $n_{614,det}/m_{det} = 0/11 = 0$<br>$n_{614,relativistic} = 2$ (out of 12); $n_{614,rel}/m_{rel} = 2/12 = .167$<br>$n_{614,evolutionary} = 1$ (out of 32); $n_{614,evo}/m_{evo} = 1/32 = .031$<br>$n_{614,symbolic} = 0$ (out of 5); $n_{614,sym}/m_{sym} = 0/5 = 0$<br>$\sum n_{614,j}/m_j = .077 + 0 + .167 + .031 + 0 = .275$ | Materialistic frame = $.077 / .275 \cdot 100 = 28.0\%$<br>Deterministic frame = $0 / .275 \cdot 100 = 0\%$<br>Relativistic frame = $.0167 / .275 \cdot 100 = 60.6\%$<br>Evolutionary frame = $.031 / .275 \cdot 100 = 11.4\%$ |

Individual listings of publications and online mentions are reported in the Supplementary Tables 1-4. Full list of key terms can be found in Supplementary Table 5. Results are also accessible in our interactive website: https://jjmorosoli.shinyapps.io/newas/.

## Acknowledgements

This study was funded by the John Templeton Foundation (Genetics and Human Agency Project). J.J.M. was supported by a QIMR Berghofer International PhD Scholarship. F.K.B. was funded by an Australian Research Council Future Fellowship (FT150100147). L.C.C. was supported by a QIMR Berghofer Research Fellowship. S.E.M. was funded by an NHMRC Senior Research Fellowship (APP1103623). We thank Altmetric for providing their data free of charge for research purposes. We thank Marlee Dutton for assistance in constructing the text corpus.

## Author contributions

J.J.M., L.C.C, F.K.B., and S.E.M. contributed to all aspects of the paper, including study design, statistical analysis and writing and revisions.

## Additional information

The authors declare no competing financial interests. Correspondence and requests for materials should be addressed to J.J.M.

## Footnotes

1 Note: Topic modeling algorithms do not provide a label for each topic. Researchers give meaning to the categories by labeling them based on most common words within each topic. In the case of “asthma and empathy”, no other label seems appropriate to subsume those keywords.

## Notes

https://jjmorosoli.shinyapps.io/newas/

### Competing Interest Statement

The authors have declared no competing interest.

### Summary of Updates

Abstract updated to include link to web app.

